# Evolution of horn length and lifting strength in the Japanese rhinoceros beetle *Trypoxylus dichotomus*

**DOI:** 10.1101/2023.02.16.528888

**Authors:** Jesse N. Weber, Wataru Kojima, Romain Boisseau, Teruyuki Niimi, Shinichi Morita, Shuji Shigenobu, Hiroki Gotoh, Kunio Araya, Chung-Ping Lin, Camille Thomas-Bulle, Cerisse E. Allen, Wenfei Tong, Laura Corley Lavine, Brook O. Swanson, Douglas J. Emlen

**Affiliations:** Department of Integrative Biology, University of Wisconsin-Madison, Madison, WI USA; Graduate School of Sciences and Technology for Innovation, 1677-1 Yoshida, Yamaguchi University, Yamaguchi 753-8511, Japan; Division of Biological Sciences, The University of Montana, Missoula MT, USA; Division of Evolutionary Developmental Biology, National Institute for Basic Biology, 38 Nishigonaka Myodaiji, Okazaki 444-8585, Japan; Trans-Scale Biology Center, National Institute for Basic Biology, 38 Nishigonaka Myodaiji, Okazaki 444-8585, Japan; Department of Science, Graduate School of Integrated Science and Technology, Shizuoka University, Shizuoka Japan; Graduate School of Social and Cultural Studies, Kyushu University, 4-2-1 Ropponmatsu, Chuo, Fukuoka 810-8560 Japan; Department of Life Science, National Taiwan University, No.88, Sec. 4, Tingzhou Rd., Taipei 11677, Taiwan; Cornell Laboratory of Ornithology, Ithaca NY; Department of Entomology, Washington State University, Pullman, Washington 99164; Department of Biology, Gonzaga University, 502 East Boone Avenue Spokane, WA 99258-0102

**Keywords:** Sexual selection, male competition, animal weapons, phylogeography

## Abstract

Rhinoceros beetle (*Trypoxylus dichotomus*) males have pitchfork-shaped head horns, which they use to pry rival males from the trunks of trees. In the largest males these horns can be three times the length of horns in the two closest sister species. Because this weapon functions as a lever, longer horns should lift with less force than shorter horns (the ‘paradox of the weakening combatant’) unless other elements of the weapon system (e.g., input lever length, muscle mass) evolve to compensate. We used next-generation sequencing approaches to consolidate 23 sample locations into 8 genetically distinguishable populations, reconstructing their historical relationships and providing a comprehensive picture of the evolution of this horn lever system. We show that head horns likely increased in length independently in the Northern and Southern lineages. In both instances this resulted in weaker lifting forces, but this mechanical disadvantage was later ameliorated, to some extent and in some locations, by subsequent reductions to horn length, changes in muscle size, or by an increase in input lever length (head height). Our results reveal an exciting geographic mosaic of differences in weapon size, weapon force, and in the extent and nature of mechanical compensation. Reconstructing the evolution of this weapon system offers critical insights towards meaningfully linking mating system dynamics, selection patterns, and diversity in sexually selected traits.

## Introduction

Sexual selection can lead to rapid increases in male weapon size (Darwin 1871; Andersson 1994). Because these weapons are deployed against the weapons of conspecific rivals, an aspect of the social environment that is itself evolving, male competition can generate consistent and intense directional selection for elaborations to weapon form that improve contest outcome (West Eberhard 1979, 1983; Lyon and Montgomerie 2012; De Lisle et al., 2021). For many weapons, this means increases in length or overall weapon size (Clutton Brock et al., 1980; Andersson 1994; Bro-Jørgensen 2007; Emlen 2008, 2014; Plard et al., 2011).

Longer weapons permit a male to touch, strike, grab or flip an opponent before that rival can do the same (e.g., Hyatt and Salmon 1978; Hongo 2003; Goyens et al., 2015; Kojima and Lin 2017; Fea and Holwell 2018). Longer weapons may also function as agonistic signals – deterrents – settling contests before they escalate into dangerous battle (Clutton Brock et al., 1979; Enquist and Leimar 1983, 1987, 1990; Maynard Smith and Parker 1976; Hyatt and Salmon 1978; Jennions and Backwell 1996; Panhuis and Wilkinson 1999; Kelly 2006; Egge et al., 2011; Biernaskie et al., 2014; Painting and Holwell 2014; Goyens et al., 2015). When opponents are evenly matched, however, even these contests escalate, and this means that weapons cannot solely serve as signals; they must also, at least on occasion, function as tools of battle (Berglund et al., 1996; Hardy and Briffa 2013; Dennenmoser and Christy 2013; McCullough et al., 2016).

While longer weapons may be advantageous for their added reach and as deterrents, longer is not necessarily better for generating force (Levinton and Allen 2005; Dennenmoser and Christy 2013; O’Brien and Boisseau 2018; Levinton and Arena 2021). Weapons that lift, pry, or squeeze function as lever systems and include an “output” lever (e.g., a horn), an “input” lever (e.g., a rigid head), and muscles attached to the input lever that, when contracted, rotate both levers about a fulcrum (Figure 1; Vogel 2013).

**Figure 1.**
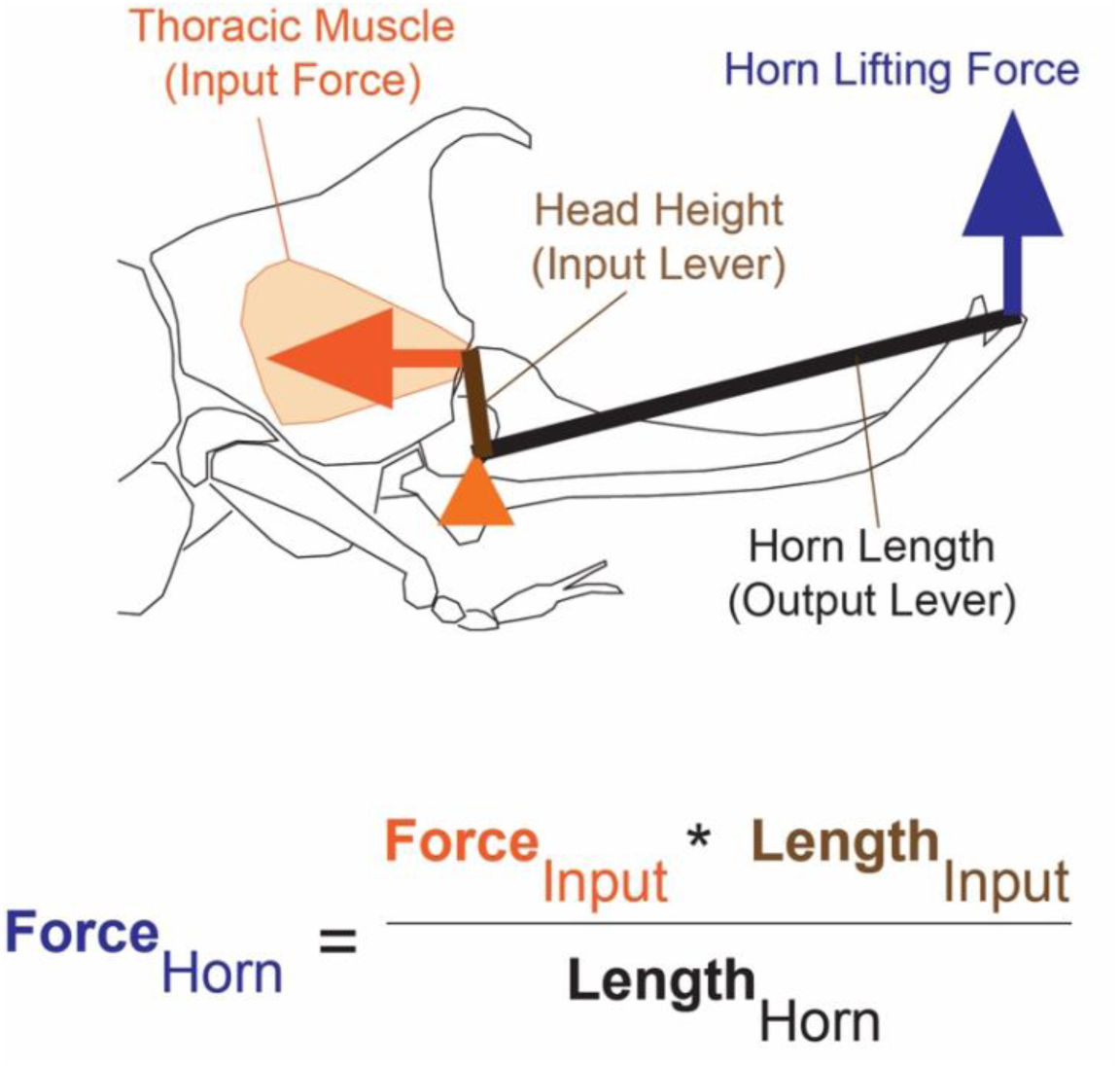
Horn lever system of *Trypoxylus dichotomus*. Males attempt to insert their horn underneath an opponent and lift them off the sides of trees. Lifting forces (blue arrow) are generated by a lever system comprising an output lever (horn length), and an input lever (head height) and muscles housed in the thorax that rotate the head (input force, orange arrow) to lift the horn (lifting force). The fulcrum (orange triangle) is located in the head between the input and output levers.

The force generated by the weapon can be predicted from the relative lengths of the two levers and the size of the associated muscles:

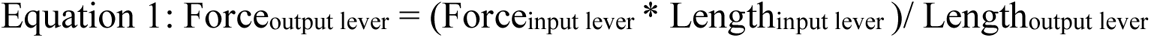

For most animal weapons the output levers (i.e., horns, tusks, antlers) are not physically constrained from expanding, protruding as they do from the body. The input levers and associated musculature, in contrast, are housed inside the body where limitations of space may constrain their size (O’Brien and Boisseau 2018). Strong sexual selection driving rapid increases in weapon length can create an imbalance with the rest of the lever system, yielding tools that lift, pry, or squeeze with reduced force (Dennenmoser and Christy 2013; Goyens et al., 2014, 2016; Mills et al., 2016; O’Brien and Boisseau 2018).

In principle, this ‘paradox of the weakening combatant’ (Levinton and Allen 2005) could limit the elaboration of sexually selected weapons, if increased weapon length reduces force enough to offset signaling or other benefits. However, compensatory changes to the lever generally maintain biomechanical force, even for the longest-weaponed individuals (see Levinton and Arena 2021 for an exception). Studies of the biomechanical force generation of animal weapons generally find that forces are maintained even in the longest-weaponed individuals due to the presence of compensatory changes to the lever system. In crabs, frog-legged beetles, stag beetles, and leaf-footed bugs increases in relative muscle size maintain weapon performance as weapon lengths increase (Levinton and Judge 1993; Goyens et al., 2014; O’Brien and Boisseau 2018; O’Brien et al., 2019); and in stag beetles and frog legged beetles the relative length of input levers increases as well (Goyens et al., 2014). Fiddler crabs and stag beetles possess an enlarged ‘tooth’ near the fulcrum, which is used to squeeze opponents close to the fulcrum where forces are strongest, effectively uncoupling the signal and weapon functions (Dennenmoser and Christy 2013; Mills et al., 2016).

To date, most biomechanical studies focus on individual variation within populations, clearly demonstrating a variety of means to mechanical compensation (e.g., static allometry of input and output levers and relative muscle size; Burrows and Wolf 2002; Westneat 2003, 2004; Waltzek and Wainwright 2003; Levinton and Allen 2005; McHenry 2011; Sutton and Burrows 2011; Dennenmoser and Christy 2013; Goyens et al., 2014, 2016; McCullough 2014; O’Brien and Boisseau 2018). A few studies extend to multiple populations or species (Swanson et al., 2013; Goyens et al., 2016; Mills et al., 2016; Buchalski et al., 2019; Palaoro et al., 2020) but none so far include sufficient information about the historical relationships of populations to permit a full reconstruction of the dynamic stages of weapon evolution and the accompanying origin and spread of mechanical compensation.

Here we present results from a comprehensive phylogeographic study of a large-weaponed species, the Japanese rhinoceros beetle *Trypoxylus dichotomus* (Linnaeus 1771; Coleoptera, Scarabaeidae). We collected samples from 23 locations across the range of this species and used high throughput DNA sequencing approaches to consolidate these into 9 genetically distinguishable populations. Using the closest sister species (*Xyloscaptes davidis*) as an outgroup, we reconstruct the historical relatedness among these populations and use this tree to examine the evolution of both weapon size, and weapon lifting strength.

We show that the most parsimonious model of horn evolution involves initial increases in weapon size associated with significant reductions to lifting strength, but this mechanical disadvantage was later ameliorated, to some extent and in some locations, either by subsequent reductions to horn length, or by an increase in input lever length (head height). In addition, some populations differ in the amount of muscle powering the horn lifting system, suggesting another mode of compensation. Our results reveal an exciting geographic mosaic of differences in weapon size, weapon force, and in the extent and nature of mechanical compensation, highlighting the ability to use extant variation among populations and their historical relatedness to support multiple distinct models for the origin and use of extreme weapons.

## Background

*T. dichotomus* is a univoltine scarab whose larvae feed in the soil on decaying wood, emerging as adults in early June to mid-July depending on the location (Plaistow et al., 2005; Iguchi 2006; Hongo 2007; Kojima et al., 2020; del Sol et al., 2020). Adult behavior has been studied most extensively on Honshu Island, Japan, where beetles fly to wounds on the sides of mature oak, ash, and maple trees (e.g., *Quercus mongolica*, *Q. acutissima*, *Q. serrata*, *Fraxinus griffithii, Acer plantanoides*; Hongo 2007) and feed on oozing sap (Siva-Jothy 1987; Setsuda et al., 1999; Hongo 2003, 2007). Males battle with rival males for ownership of feeding territories and the largest males with the longest horns are most likely to win (Obata & Hidaka 1983; Siva-Jothy 1987; Hongo 2003, 2007; Karino et al., 2005; del Sol et al., 2020). Females fly to these territories to feed and mate, before leaving to lay eggs in decomposing litter up to a kilometer or more away (McCullough et al., 2012; McCullough 2013).

In the first comprehensive field study of this species, Hongo (2003, 2007) showed that in Kyoto, Japan, a population with long male horns, feeding territories were severely limiting and mating success was confined to the largest and longest-horned males, consistent with a resource defense mating system and strong sexual selection for enlarged/exaggerated male weapons. Recently, del Sol et al., (2020) extended this study to include measures of fighting and mating success at one additional long-horned population (Kameoka, Japan) and three populations with relatively shorter horns (Yakushima Island, Japan; Chia yi and Puli, Taiwan), to test for local differences in breeding ecology and the strength of selection acting on horns. Males used horns in battles for sap-sites in all the studied locations, and in all places males with the longest horns were the most likely to win (del Sol et al., 2020). However, populations differed in the relative scarcity of feeding territories and, accordingly, in the relative probability that any particular defended territory would be visited by a female. The result was that victorious males derived greater reproductive benefits in long-horned populations than they did in the shorter-horned populations, compelling preliminary evidence that local differences in the details of the mating system may help explain population differences in relative weapon size (Hongo 2006; del Sol et al., 2020).

*T. dichotomus* populations differ in horn strength (lifting force) as well as horn length (Buchalski et al., 2019). Horns are used to pry rivals off of the sides of trees, flipping them to the ground. Specifically, the pitchfork-shaped head horn functions as a simple lifting lever, with the junction of the head and thorax serving as a fulcrum. The head acts as the input lever and is attached to large muscles housed in the thorax, which rotate the head to raise the horn (Buchalski et al., 2019). Rapid evolutionary increases in output lever (horn) length may have outpaced correlated changes in the input lever, confined as it is inside the small head, resulting in a reduction to the lifting force generated by this lever system. If true, then *T. dichotomus* horns may have suffered from the ‘paradox of the weakening combatant’ and experienced selection for mechanisms that ameliorate the mechanical disadvantage of long horns (Levinton and Allen 2005). Preliminary studies demonstrate that within populations, large males compensate for their long horns with relatively larger thoracic lifting muscles than smaller males (Buchalski et al., 2019), but across populations the pattern is less clear, motivating the present study.

## Methods and Materials

### Study system

Japanese rhinoceros beetles (‘Yamato kabutomushi’) are large, abundant, and often observed scarabs native to Eastern Asia, including mainland China, Taiwan, Korea and Japan (Laurent 2000; New 2005; Brock 2006; Kawahara 2007; Takada 2013; Wang et al., 2022). Males have a short, curved horn extending from the dorsal prothorax and a much longer, four-tined “pitchfork” shaped horn extending from the top of the head (Figure 2). Males of the two closest sister species, *Xyloscaptes davidis* and *Allomyrina pfeifferi* (Rowland and Miller 2021; Dutrillaux et al., 2013; Jin et al., 2016; Yang et al., 2020), have a much smaller, two-tined forked horn extending from the center of the head. Both *X. davidis* and *A. pfeifferi* are rare and very poorly understood outside of systematics.

**Figure 2.**
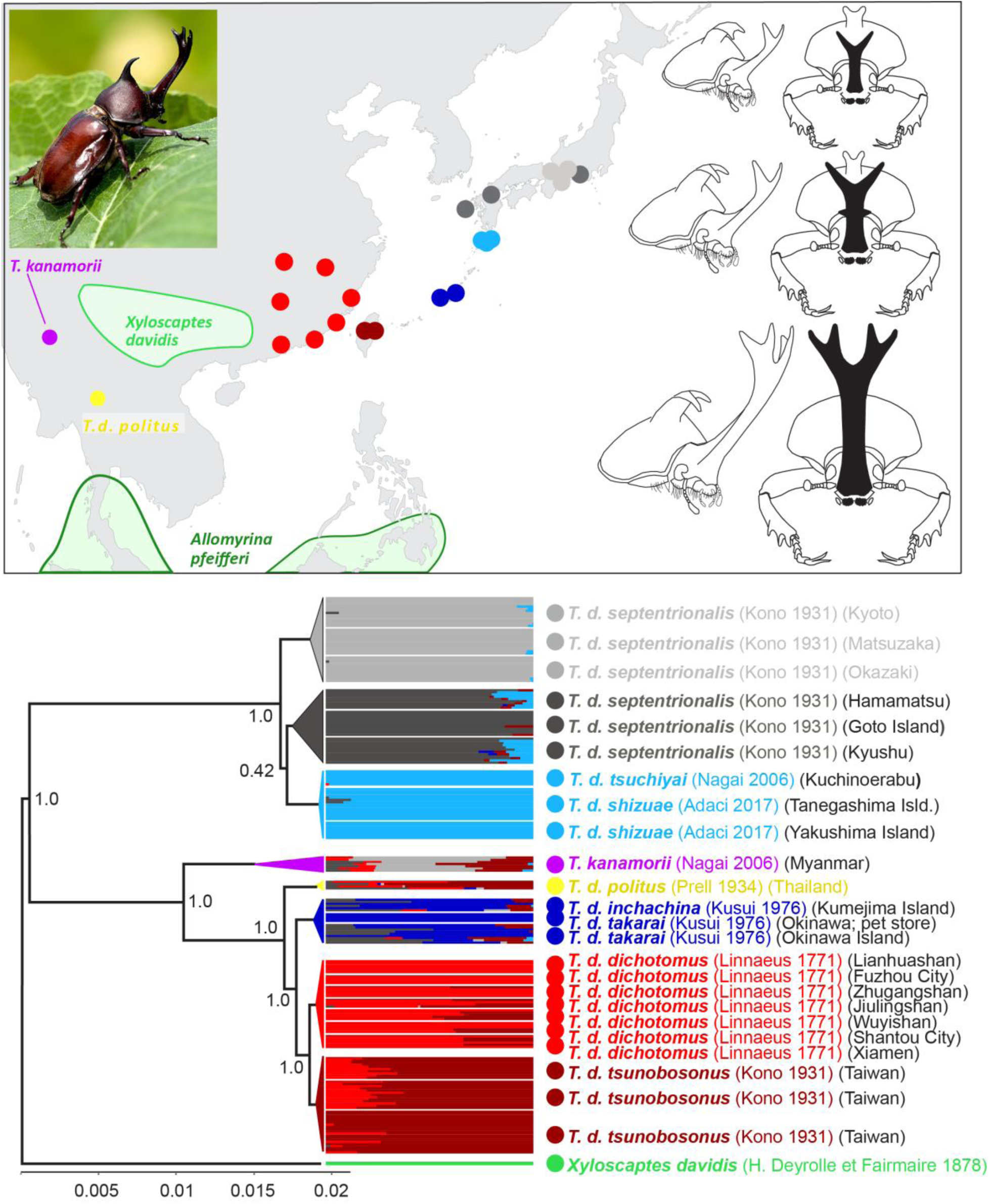
Top. Sample locations and head horns of *Trypoxylus dichotomus* (bottom), *Xyloscaptes davidis* (middle), and *Allomyrina pfeifferi* (top). Bottom. Ancestral relationships among populations of *T. dichotomus*. Genetic structure analyses show that individuals cluster into 6 primary populations: Japan Central (light gray), Japan South (dark gray), Kuchinoerabu/Yakushima/Tanegashima islands (light blue), Okinawa/Kumejima (dark blue), mainland China (red) and Taiwan (maroon). *T. kanamorii* (purple) and *T. d. politus* (yellow) were also considered to be separate populations based on morphology and geographic location. Goto island beetles, although placed within the Japan South cluster, also were morphologically distinct, and thus were treated as a separate population for horn length and horn strength comparisons. Clade support values represent Bayesian posterior probabilities.

Ten subspecies of *T. dichotomus* have been described based on geographic location and morphology, including conspicuous and consistent population differences in relative horn length (*T. d. septentrionalis* [Korean Peninsula, Japan; Kono 1931; Nagai 2007]; *T. d. tsuchiyai* [Kuchinoerabu Island; Nagai 2006]; *T. d. shizuae* [Yakushima and Tanegashima Islands; Adachi 2017]; *T. d. dichotomus* [Mainland China; Linnaeus 1771, Nagai 2007]; *T. d. tsunobosonus* [Taiwan; Kono 1931; Nagai 2007]; *T. d. takarai* [Okinawa; Kusui 1976]; *T. d. inchachina* [Kumejima; Kusui 1976]; *T. d. politus* [Myanmar, Thailand, Vietnam; Prell 1934; Nagai 2007]; *T. d. shennongjii* [Shennongji, Hubei, China; Satoru 2014]; and *T. kanamorii* [Myanmar; Nagai 2006]).

### Sample collection and assignment to genetic populations

Our approach comprised the following steps, each described in detail below: (1) samples were collected from 23 locations across the range of the species and including 9 of the 10 described subspecies (Table S1); (2) genomic DNA was extracted from approximately 10 animals per location (198 beetles total) and sequenced to identify approximately 13,500 informative SNPs distributed across the genome; (3) we assigned animals from the different locations to 8 genetically distinguishable populations; (4) two individuals from each of these populations and from the outgroup *X. davidis* were then used to perform coalescent analyses of population ancestry, generating a well-supported phylogeny for the historical relationships among the populations; animals from all of the sample localities were then pooled into their respective genetic populations and these groupings were used for morphological comparisons of relative horn length (4) and for biomechanical calculations of horn strength (5).

Once the genetic populations had been defined, we added additional samples (n = 1375) from these same regions to increase our sample sizes for the horn length versus body size allometry estimation, and to improve our ability to test for differences in relative horn length across populations. Because these additional animals were from private collections we did not use them for the biomechanical force estimation, which would have required destructive sampling of the specimens.

### Population Genetic Analyses

#### DNA extraction and ddRAD library preparation

For DNA extraction, we removed cuticle and isolated ∼20mg of muscle tissue from each individual sample, then performed tissue lysis and DNA isolation using the Wizard SV Genomic DNA Purification kit (Promega). We fluorometrically quantified DNA using the PicoGreen dsDNA kit (Life Technologies) and constructed double digest restriction-site associated DNA sequencing libraries for Illumina Sequencing following Peterson et al., (2012). Briefly, after normalizing samples to 15, 30 or 90 ng, we performed double restriction digests using the enzymes NlaIII and MluCI (New England Biolabs). All enzymatic steps were followed by cleanups with solid-phase reversible immobilization beads (Sera-mag Speedbeeds, Fisher). Post digest, we performed ligations with adapters P1 and P2, pooled samples (<48 samples per pool), and ran each pool on one lane of a Pippen Prep 2% MarkerB 100–600 bp cassette (Sage Science) to collect fragments between 426-462bp in length, which includes additional adapter sequences. We then enriched for fragments ligated to P2 adapters using streptavidin-coupled beads (Dynabeads M-270, Invitrogen). Finally, we amplified each pool using 4 separate PCRs, each with 12-cycles (Phusion High Fidelity PCR kit, New England Biolabs).

#### Mapping and genotyping

All libraries were sequenced using 2×150bp reads on an Illumina HiSeq 4000 machine. After sequencing, we demultiplexed raw reads by individuals using the “process_radtags” function in Stacks version 1.13 (Catchen et al., 2013; Rochette et al., 2019). We next removed PCR duplicates (facilitated by a unique molecular index located on the i7 index read and a custom python script) and mapped reads to a draft *T. dichotomus* genome (Morita et al., 2022) using BWA-MEM (Li 2013). We calculated genotype probabilities using the mpileup algorithm (options: flags -C 50, -E, -S, -D, -u, -I) and performed genotype calls using BCFtools, with both functions available in the SAMtools package (Version 1.5; (Li et al., 2009; Li 2011)).

After examining several measures of read depth and quality, we implemented an initial filter to retain only genotypes with a minimum per individual depth of 8 and minimum genotype quality of 30, and only kept sites where the mean depth was greater than 1. We next removed any sites where less than 50% of individuals had genotype calls. We further removed 500bp windows (i.e., full RADtags) containing more than six SNPs with high depth reads (>1000), as well as any regions containing SNPs with total depth >1300 (((screen_sites_depth_to_drop.R script—output =South_phylo_6pop_011018_minIndDP8_Max6snpsPerCluster_maxDP1300.nex)). Additional filters specific to different analyses are noted when appropriate in the following sections.

#### Population genetic analyses

We used a cross validation approach implemented in the software package ADMIXTURE (Alexander et al., 2009) to estimate the number of population clusters. Because high LD can lead to spurious signals, we first thinned our dataset by identifying any SNPs located within 50bp of each other and having an R^2^ value > 0.1, and then only keeping the site with the highest minor allele frequency. For the ADMIXTURE runs, we tested subpopulation (k) values ranging from four to nine, and determined the best model fit by identifying the k value that minimized CV error.

### Coalescent Analyses and Phylogeny Construction

We estimated phylogeny using beetles collected from 11 of our 23 sample locations, representing all of the genetically distinguishable populations identified by the ADMIXTURE analyses and all 9 of the formally described subspecies included in this study (Table 1). Specifically, we compiled genotype data for two beetles from each of the 11 sample locations, keeping individuals with the most complete genotype coverage in each population, as well as one outgroup individual from the sister species *Xyloscaptes davidis*. We then estimated population ancestry using a full coalescent analysis (SNAPP; Bryant et al., 2012) implemented in BEAST v2.4.5 (Bouckaert et al., 2014). For the SNAPP analysis, we used a gamma rate prior, used the built-in function to estimate mutation rates U and V, and ran the MCMC for up to 2 million cycles, storing data every 1,000 cycles. All other parameters were run with default settings. Runs were continued until the estimated sample size (ESS) for the posterior probabilities exceeded a value of 200. ESS’s for other model parameters were also evaluated with the Tracer v1.7 software (Rambaut et al., 2018).

We first ran a full tree using all of the genetically distinguishable *T. dichotomus* populations (eleven of the sample locations) and *X. davidis* as an outgroup. This tree revealed a well-supported deep split between northern and southern *Trypoxylus* lineages, suggesting the beetles colonized these regions independently with each clade having its own evolutionary trajectories. However, our DNA from *X. davidis* was degraded relative to our other samples, reducing the number of usable SNPs. We therefore ran separate, clade-specific analyses for these northern and southern trees, using a representative from the other clade as an outgroup instead of *X. davidis*, increasing the number of usable SNPs to 747 and 4,091 respectively.

The northern analysis included a total of 6 locations (Kyushu, Kyoto, Matsuzaka, Hamamatsu, Yakushima Island and Tanegashima Island) and Wuyishan, China, as an outgroup, while the southern analysis used 5 locations (*T. kanamorii* [Myanmar]; *T. d. politus* [Thailand]; Okinawa, Fuzhou City, and Taiwan) and Matsuzaka, Japan, as an outgroup. Finally, we sampled from the resulting tree files, after removing the first 10% of traces as a burn-in, to estimate branch lengths and generate posterior probabilities for all nodes. Tree annotation and visualization was performed using the Densitree (Bouckaertand Heled 2014) and treeanotator software supplied with BEAST2, as well as FigTree (Rambaut 2017).

### Weapon Size Comparisons

As in other insect taxa with highly plastic sexually selected structures (e.g., fly eyestalks, Wilkinson 1993, David et al., 1998, Knell et al., 1999, Baker et al., 2001; earwig forceps, Tomkins 1999, Tomkins & Brown 2004; dung beetle horns, Emlen 1994, 1996; Moczek & Emlen 1999; flour beetle mandibles, Okada & Miyatake 2009, 2010; frog-legged leaf beetle hindlegs, O’Brien et al., 2017), *Trypoxylus* horn length *per se* has little detectable heritability in contemporary populations (Karino et al., 2004; Johns et al., 2012) and horn length evolution likely arises through changes in the developmental mechanisms modulating horn growth in relation to the nutritional state and overall body size of an animal (see, e.g., Ito et al., 2013; Emlen et al., 2012; Gotoh et al., 2015; Ohde et al., 2018; Morita et al., 2019; Adachi et al., 2020). Therefore, we compare relative sizes of horns across populations by comparing the steepness, elevation, and/or shape of population-level patterns of horn length / body size scaling (Emlen & Nijhout 2000; Emlen & Allen 2004; Shingleton et al., 2007; Dreyer et al., 2016; O’Brien et al., 2017).

All statistical analyses were performed in R (version 4.1.1; R core team 2021). For all models assuming normally distributed and homoscedastic errors, we systematically checked and confirmed this assumption by plotting the residuals against normal quantiles and residual vs fitted values. We first compared the horn length / body size scaling relationships between the outgroup *X. davidis* and the northern and southern *T. dichotomus* clades. We fitted a linear mixed model (function lme, R package “nlme” (Pinheiro et al., 2021)) including log10 horn length as the response variable and log10 thorax width, population, and their interaction as fixed effects.

The population of origin was included as a random effect. Significance of the fixed effects was assessed using a type I (sequential) ANCOVA. This allowed us to test for differences between major clades in relative horn length, and to attribute any differences to changes in horn allometry slope and/or intercept. Post-hoc pairwise intercept and slope comparisons were performed using estimated marginal means and Tukey contrasts (functions emmeans or emtrends in R package “emmeans”).

Next, we compared *T. dichotomus* populations within the northern and southern clades separately. However, in contrast to *X. davidis*, horn-length body size scaling relationships in *T. dichotomus* are not linear, even when log-transformed (Iguchi 2002; Hongo 2007; Knell 2009; McCullough et al., 2015; Packard 2021; del Sol et al., 2020), so we also compared *T. dichotomus* populations by fitting the sigmoid equation:

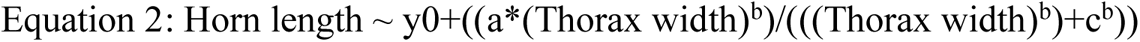

Where y0 is the minimum horn length, a is the horn length range, b is the curve steepness, and c is the body size (thorax width) at the inflection (Parzer and Moczek 2008; Casasa and Moczek 2018). For both the northern and southern clade, we went through a model selection procedure to identify which parameters were better considered to be population-specific. We started by fitting the full model with all four parameters defined as population-specific and compared with four nested models each fixing one of the parameters as shared across populations. The model with the lowest Akaike information criterion (AIC) was selected and the procedure repeated until no simpler model was found. Using the best selected model, we tested for pairwise population differences for the parameters fitted as population specific using the Bonferroni method to account for multiple testing (function pairComp in package “aomisc”; Onofri 2020). This permitted us to focus specific contrasts around the changes in horn length inferred from the phylogeny (i.e., not simply compare all populations to each other but compare before and after populations that bracket each inferred transition), and to describe more precisely how horn allometries evolved.

### Evolution of the horn lever system: head height, thoracic muscle area, and estimates of horn lifting force

Morphological variables were recorded from beetle specimens collected in the field and frozen or dried prior to transport. Measurements were made using digital calipers to the nearest 0.01 mm. Output lever of the horn system was the distance from the fulcrum hinge point at the back of the head to the end of the longest tine of the horn. Input lever of the horn system was the distance from the fulcrum to the insertion of the dorsal prothoracic muscle at the dorsal/caudal peak of the head. Prothorax width was measured at the widest point of the prothorax. Because we were dealing with frozen or dried specimens, muscle cross sectional area was estimated from the area of the sclerotized plate where the dorsal prothoracic muscle originates at the caudal end of the prothorax. Beetles were dissected, digital images were recorded of this plate, and area was calculated with ImageJ (Schneider et al., 2012).

Muscle force per cross sectional area was estimated by measuring lifting forces from live beetles (see Buchalski et al, 2019 for details). Therefore, our predictions of horn lifting force reflect changes in mechanical advantage and muscle cross sectional area across populations and do not include possible differences in muscle physiology.

Horn lifting forces were predicted using Equation 1 with the following measures:

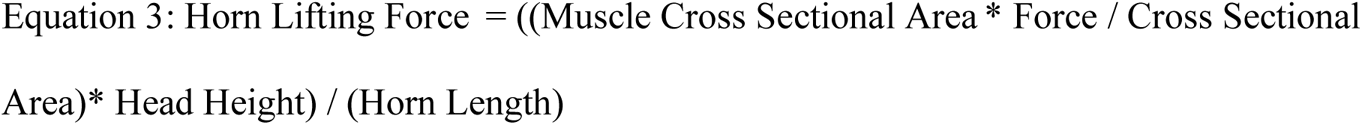

Head height (input lever length), muscle cross-sectional area (input force), and horn lifting forces were first compared between the outgroup *X. davidis* and the two major *T. dichotomus* clades (i.e., northern and southern). For each biomechanical variable, we fitted a linear mixed model including log10-transformed head height, muscle cross-sectional area, or horn lifting forces as the response variable and log10 thorax width, population, and their interaction as fixed effects (function lme, R package “nlme”). The population of origin was included as a random effect. Significance of the fixed effects was assessed using a type I (sequential) ANCOVA. Non-significant interactions (p > 0.1) were removed from final analyses. Post-hoc pairwise intercept and slope comparisons were performed using estimated marginal means and Tukey contrasts (functions emmeans or emtrends in R package “emmeans”).

We then compared *T. dichotomus* populations within the northern and southern clades separately. Log10-transformed head height, muscle cross-sectional area, or horn lifting force was included as a response variable in a linear model, with log10-transformed prothorax width and population, and their interaction (if significant, i.e., p<0.1) included as explanatory variables. Post-hoc pairwise comparisons between populations (intercept and slope if the interaction was significant) were performed using estimated marginal means and Tukey contrasts (functions emmeans or emtrends in R package “emmeans”).

## Results

### Genetic structure of T. dichotomus populations

After filtering, our population structure dataset contained 198 individuals genotyped across 13,547 sites. ADMIXTURE analyses suggest that the genetic variation in *T. dichotomus* is best explained (CV error = 0.22752) by grouping individuals into the following six general subpopulations: Japan central; Japan south, including Goto Island; Kuchinoerabu-Yakushima-Tanegashima Islands; Okinawa-Kumejima Islands; Taiwan; and mainland China (Figure 2). k=5 populations also provided a good fit (CV error = 0.22828), with the main difference being that the small islands of Kuchinoerabu, Yakushima, and Tanegashima, which lie off the southern coast of the larger island of Fukuoka, Japan, cluster with one of the Japanese main island populations.

Beetles from *T. kanamorii* and *T. d. politus* each showed a mixture of genetic backgrounds; however, it is unclear whether this pattern results from vicariance followed by incomplete lineage sorting, or by recent gene flow between divergent lineages. Because these subspecies are clearly distinguishable morphologically, and both are rare and confined to the far western part of the range (Myanmar and Thailand), we treated them as separate populations in this study.

Beetles on Goto Island cluster with the southern Japanese main island population and were not distinguishable in our admixture analysis. However, males on Goto Island are morphologically distinct from mainland beetles (relatively shorter horn lengths; Figure 3), so we treated Goto as a separate population for horn length and horn strength analyses. Consequently, we assigned beetles into nine genetically and/or morphologically recognizable populations (Table S1, Figure 2) and used these groupings of individuals for the subsequent horn length and horn strength analyses.

**Figure 3.**
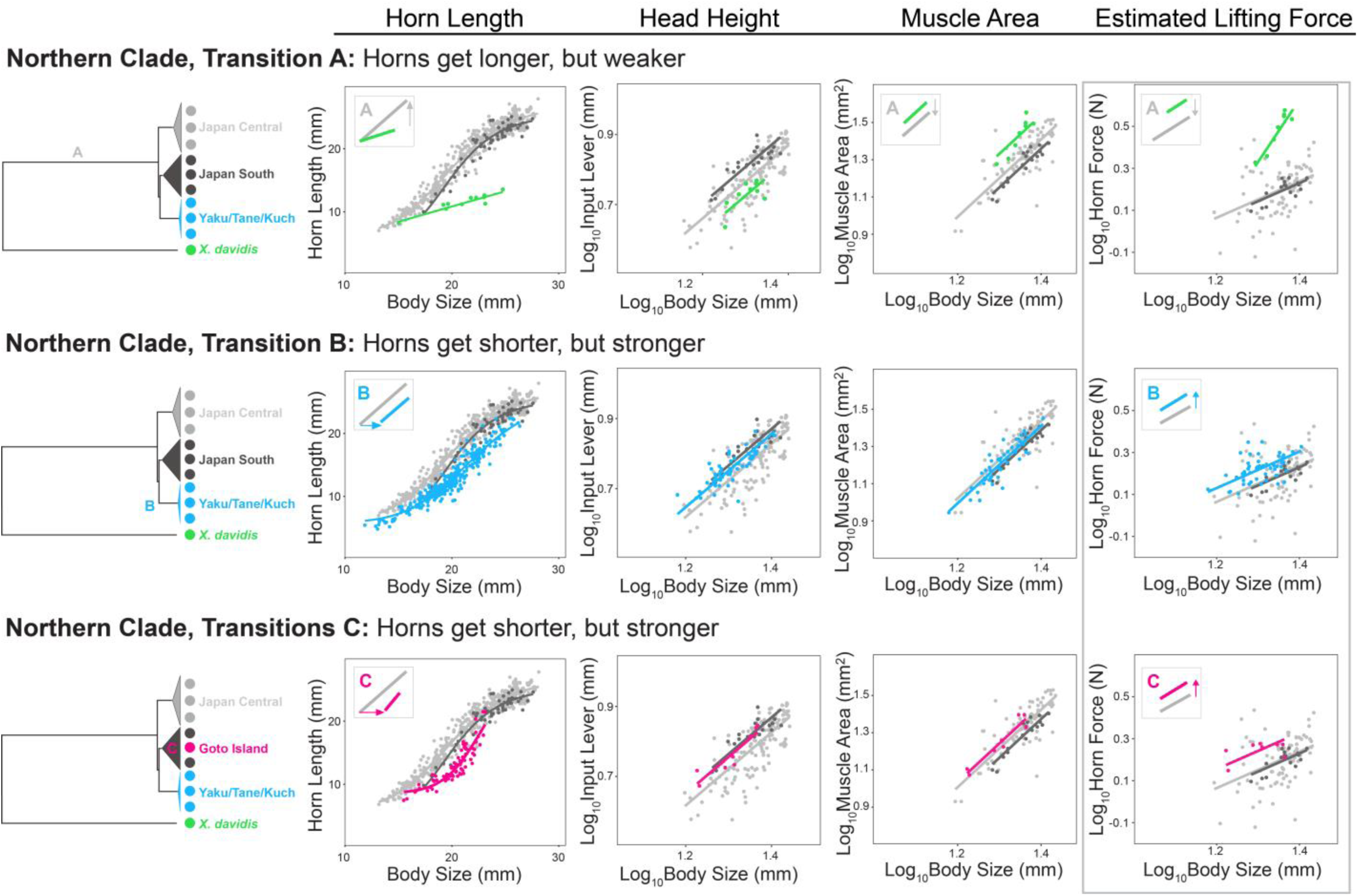
Inferred transitions in horn length, head height (input lever length), muscle area, and estimated horn lifting force in the northern lineage of *T. dichotomus*. Horns of *X. davidis* (green) were considered to represent the ancestral morphology of this species. Significant population differences are indicated by insets. Analyses are presented in Tables S3-S13.

### Historical phylogeography of T. dichotomus

Both the combined analysis with *X. davidis* as an outgroup, and the separate northern and southern clade analyses, each with a representative of the other clade as outgroup, yielded the same tree topology, and all but one node had posterior probabilities of 1.0 (Figure 2). Our results clearly support a deep early split between northern and southern populations of beetles, and this split appears to pre-date the branching of *T. kanamorii* and *T. d. politus*.

The northern lineage includes two clusters on the Japanese main islands (*Trypoxylus dichotomus septentrionalis*), and a cluster that includes three of the tiny offshore islands, Kuchinoerabu (*T. d. tsuchiyai*), and Yakushima and Tanegashima (*T. d. shizuae*) (Figure 2). The southern lineage includes *T. kanamorii* and *T. d. politus* (both sampled from Thailand), as well as Okinawa-Kumejima (*T. d. takarai* / *T. d. inchachina*), Mainland China (*T. d. dichotomus*), and Taiwan (*T. d. tsunobosonus*).

T. kanamorii and T. d. politus were predicted to be the most basal lineages in our study based both on geographic location and on morphological taxonomy (Prell 1934; Nagai 2006), and our population genetic results suggest they each contain mixtures of alleles from both northern and southern populations. However, our coalescent analyses strongly support placement of these populations within the southern lineage, rather than at the base of the combined T. dichotomus tree. If true, then this would indicate that long horns evolved independently in the northern and southern lineages (see below).

The islands Okinawa and Kumejima also share alleles with both Northern and Southern populations, yet they too clearly branch from the southern lineage. In the case of *T. kanamorii* and *T. d. politus*, we suspect that the shared alleles may reflect genetic variation present in an ancestral population predating the north-south split. This may also be true for Okinawa-Kumejima. However, their location roughly equidistant between Japan and Taiwan suggests that secondary colonization and gene flow could also account for the mixture of north-south alleles that they contain. Additional analyses will be needed to more fully resolve these issues.

Regardless, our results provide a well-supported tree sufficient for reconstructing historical patterns of evolution of horn length and horn strength.

### Evolution of male horn length

*Trypoxylus dichotomus* males had relatively longer horns and their horn length / body size allometry had a steeper slope than males of the most closely related sister species, *Xyloscaptes davidis* (Figures 3, 4; Figure S2, Table S2).

**Figure 4.**
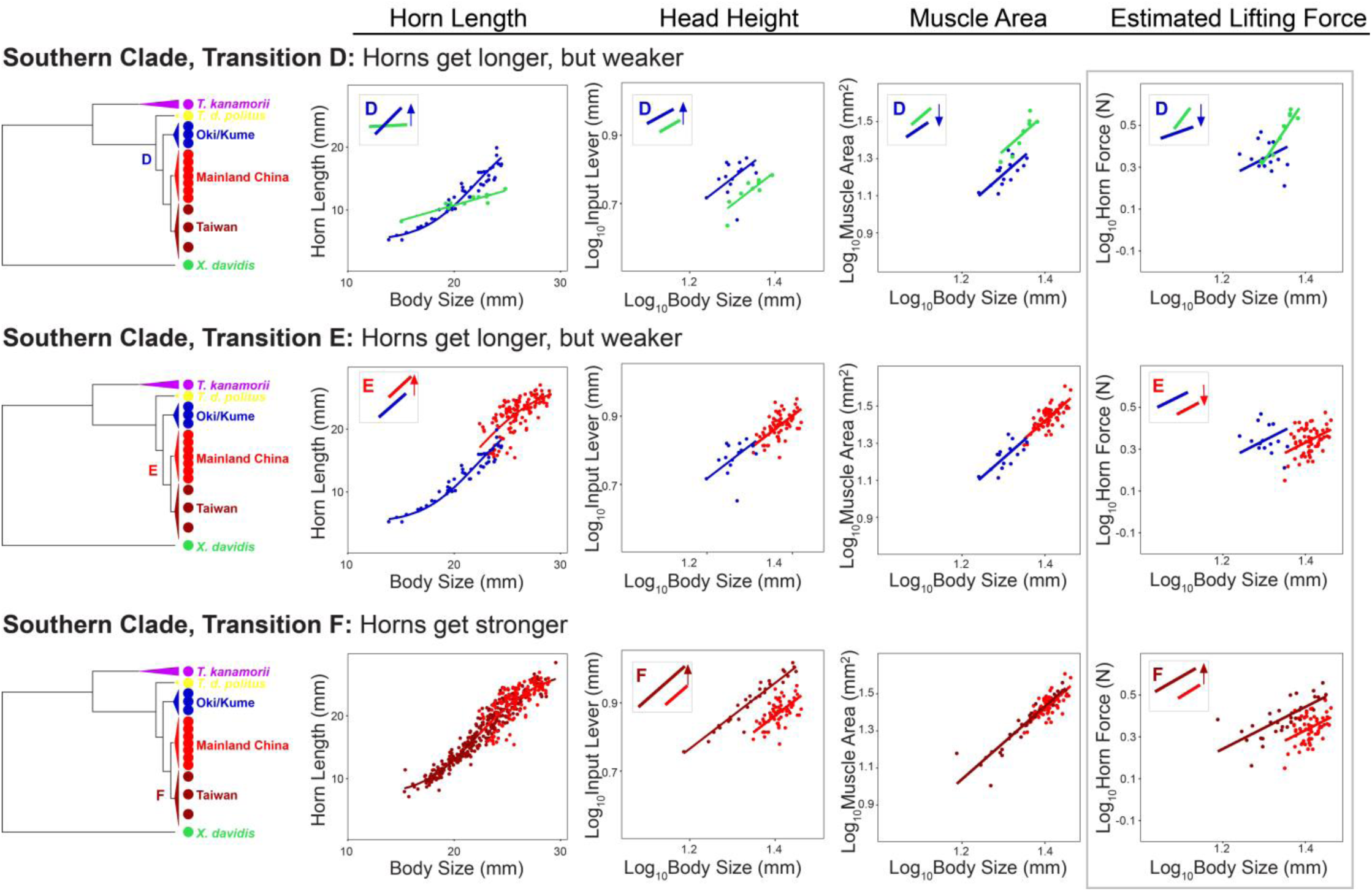
Inferred transitions in horn length, head height (input lever length), muscle area, and estimated horn lifting force in the southern lineage of *T. dichotomus*. Horns of *X. davidis* (green) were considered to represent the ancestral morphology of this species. Significant population differences are indicated by insets. Analyses are presented in Tables S3-S13.

When *Trypoxylus* populations were mapped onto the phylogeographic tree and compared using parameter substitutions to a sigmoid curve (Tables S3, S4; Figures S3, S4), horn evolution was reconstructed as follows. In the north (Figure 3), *Trypoxylus* experienced a dramatic early increase in relative horn length resulting from an increase in the slope of the horn length / body size scaling relationship (Transition A), leading to long-horned beetles in the Japanese main islands. Then, presumably as beetles colonized the tiny offshore islands of Kuchinoerabu, Yakushima, and Tanegashima, horn sizes got smaller as the allometry intercept shifted to a larger body size (Transition B). Finally, although our ADMIXTURE analyses still cluster Goto Island beetles with the Japan South population, the horn lengths of Goto males are significantly shorter than main island beetles resulting from a shift in the allometry intercept and an increase in allometry steepness (Transition C). Interestingly, although the relative horn lengths (intercepts) of Goto and Kuchinoerabu-Yakushima-Tanegashima beetles do not differ – males have relatively short horns in all of these islands – the steepness of the sigmoid scaling relationship does differ between Goto and the other offshore island populations (Figure 3C; Table S3), a morphological difference consistent with our genetic evidence which suggests a separate colonization event and an independent evolutionary reduction in relative horn length.

In the Southern clade (Figure 4), male horn length shows similar evolutionary dynamics. Both *T. kanamorii* and *T. d. politus* are rare, and we were only able to measure 3 male specimens of *T. kanamorii* for this study. However, both subspecies are known to have very short horns, and the three male *T. kanamorii* we included fell along the same horn length / body size allometry as the short-horned sister species *X. davidis* (Figure S1). From these short-horned ancestors beetles appeared to have evolved several changes in horn length. Understanding the directions and magnitude of change in horn length in this Southern lineage depends on knowing the state of the last common ancestor of Okinawa-Kumejima and mainland China animals, which is difficult to interpret given that there are too few lineages to perform a robust ancestral state calculation and recent studies show that island ecologies favor shorter horns (Nagai 2007; Adachi 2017; del Sol et al., 2021). At least one change in horn length occurred in Okinawa-Kumejima males. For simplicity, we describe this as an evolutionary increase leading to a slightly steeper allometry than *X. davidis* (Transition D). A second evolutionary change involved an increase in both allometry steepness and intercept in the lineage of long-horned males of the Mainland China and Taiwan populations (Transition E).

### Evolution of the horn lever system: head height, thoracic muscle area, and estimates of horn lifting force

Population comparisons of head height (input lever length), relative muscle size, and horn lifting force are presented in Tables S5-S13 and Figures S5-S13.

In the North, the initial increase in relative male horn length (output lever length) resulted in a significant reduction in horn lifting force (Transition A in Figure 3; Table S11). However, when beetles colonized the offshore islands Kuchinoerabu-Yakushima-Tanegashima and Goto, male horn lengths subsequently evolved to shorter lengths, leading to increases in horn lifting strength (Transitions B, C in Figure 3; Table S12).

In the South, males in Okinawa-Kumejima evolved changes in both horn length (Transition D) and head height (input lever length), so that they lift with the same relative horn strength as *X. davidis* (Figure 4). However, as horn length increased in the lineage leading to the long-horned males of the mainland China population (Transition E), head height did not change and these beetles wield weapons with significantly weaker forces than their shorter-horned ancestors.

Interestingly, males in Taiwan have the same muscle mass and horn lengths relative to body size as Mainland China males but, due to a dramatic evolutionary increase in head height, they lift with significantly stronger forces than their mainland counterparts (Transition F) (Tables S7, S13).

## Discussion

Our phylogenetic and population structure results (Figure 2) agree overall with a recent study using specific-locus amplified fragment sequencing (Yang et al., 2020). For example, Yang et al., (2020) also find a deep split between northern and southern populations; they also place *T. kanamorii* and *T. d. politus* within the southern clade, rather than basal to the *Trypoxylus* tree; and they also find Okinawa-Kumejima beetles to be genetically very distant from other populations (see also Hosoya and Araya 2010). However, Yang et al (2020) place Okinawa-Kumejima in a cluster with Taiwan beetles, whereas we find Taiwan strongly and closely related to beetles from mainland China.

The Ryukyu archipelago formed sometime between the Cretaceous and early Miocene (>12 million years ago) along the eastern rim of a contiguous terrestrial continental shelf (Nakada et al., 1991; Government of Japan 2019), and many plants (Chiang & Schaal 2006; Nakamura et al., 2009), mammals (Millien-Parra & Jager 2017), termites (Park et al., 2006), and stag beetles (Huang & Lin 2010) colonized the region at this time, before rising sea levels isolated islands in the chain. A recent fossil-calibrated molecular clock study estimated the divergence time for the split between *Xyloscaptes davidis* and *Trypoxylus dichotomus* at 29 mya (95%: 18 – 44 mya; Jin et al., 2016), so ancestral populations of *Trypoxylus* likely would have been present when the Ryukyu islands were still connected to the mainland continental shelf. We suspect *T. dichotomus* first colonized the islands at this time (Figure 5A) and were subsequently stranded when sea levels rose and cut off the Ryukyu island chain.

**Figure 5.**
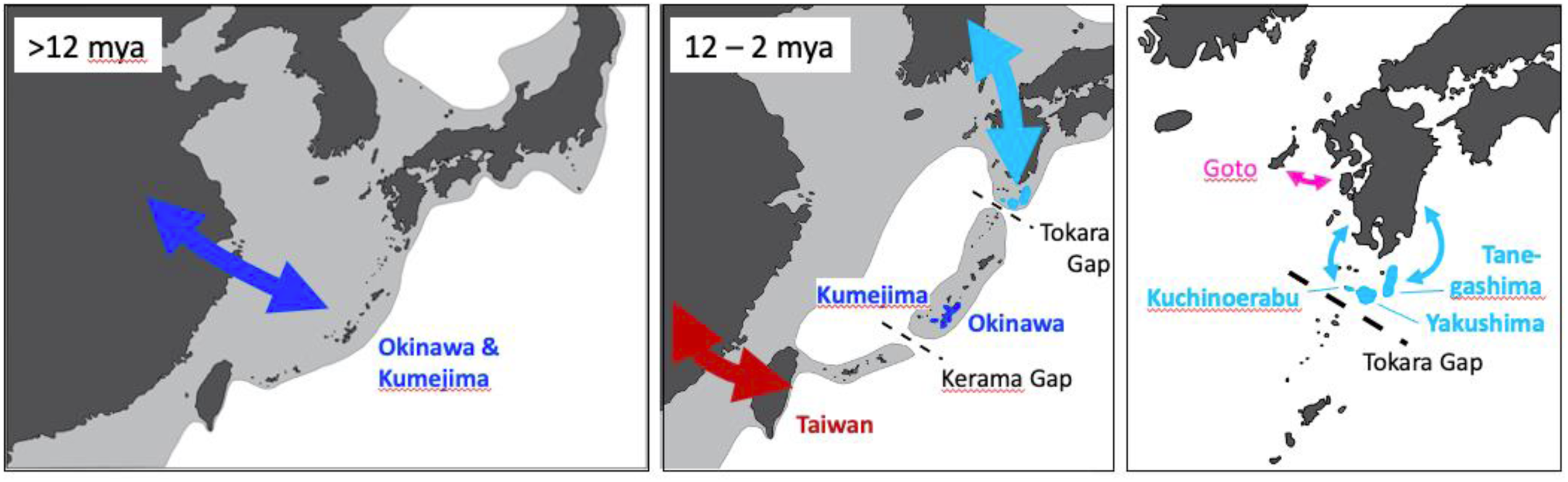
Hypothesis for the colonization history and ancestral relationships among populations of *T. dichotomus*.

Repeated drops in sea level during Pleistocene glaciation cycles temporarily re-connected islands at the northern and southern ends of the Ryukyu chain to the mainland (Figure 5B), providing episodic opportunities for mainland beetles to re-colonize some of the islands, hybridizing with or replacing existing populations. In contrast, beetles living in the middle of the archipelago (e.g., Okinawa, Kumejima), may have remained stranded due to deep oceanic trenches separating them from islands to the north and south (Tokara and Kerama Gaps, respectively; Nakada et al., 1991). This would account for both the ancestral horn morphology of these beetles (see below), and their genetic distance from beetles on islands to the north and south (Hosoya and Araya 2010). The climatic and faunal records are also more consistent with our phylogenetic result placing the Okinawa-Kumejima cluster distinct from one containing Taiwan and China, the latter of which would have had multiple opportunities to exchange migrants when the lands were connected during Pleistocene Ice Ages.

Our genetic data also suggest that colonization of the tiny islands adjacent to the Japanese main islands (Goto, Kuchinoerabu, Yakushima, Tanegashima) occurred at least twice and much more recently than colonization of Okinawa-Kumejima, either across the Pleistocene land bridges (Figure 5B) or through dispersal events from the main islands (Figure 5C). Subsequent studies of gene flow and demography will be needed to better test these hypotheses.

Horns in *T. dichotomus* almost certainly started small. Both of the sister species, *Xyloscaptes davidis* and *Allomyrina pfeifferi*, and the two most basal populations of *Trypoxylus*, *T. kanamorii* and *T. d. politus*, have short horns, and beetles isolated on the mid-Ryukyu islands of Okinawa and Kumejima have horns only slightly longer. Then, based on our genetic and phylogeographic results, male horn lengths increased dramatically two times, more than doubling in length along both the northern and southern lineages, respectively.

Both increases in horn length resulted from the evolution of steeper horn length / body size allometry slopes (Table S2; Figure 3, 4). For decades biologists have debated the extent to which morphological trait allometries constrain the evolution of animal morphology (e.g., Huxley 1924, 1932; Gould 1966, 1977; Klingenberg 2005; Voje et al., 2014; Houle et al., 2019), and the slope in particular is famously conserved (Voje et al., 2014; Pélabon et al., 2014; Houle et al., 2019).

Numerous artificial selection experiments have altered the slope of a static trait allometry (e.g., wing size in *Drosophila melanogaster* [Bolstad et al., 2015; Stillwell et al., 2016] and the butterfly *Bicyclus anynana* [Frankino et al., 2005]; and eyestalk length in the fly *Cyrtodiopsis dalmanni* [Wilkinson 1993]) and developmental genetic studies now point towards candidate genes and physiological pathways that could contribute to static allometry slope evolution (Shingleton et al., 2007; Shingleton and Mirth 2008; Tang et al., 2011; Emlen et al., 2012; Shingleton and Tang 2012; Gotoh et al., 2015; Cooper 2019; Okada et al., 2019; Toubiana et al., 2021; Vea and Shingleton 2021). Yet, it is also clear that changing a static allometry slope is not easy – responses to selection are erratic and much slower than responses to selection applied to the intercept of these same allometries (e.g., Bolstad et al., 2015; Stillwell et al., 2016). Indeed, a meta-analysis of more than 300 empirical studies of static trait allometry evolution concluded that allometry *slopes* likely change slowly over long time scales (> 1 million years) in contrast with allometry *intercepts*, which routinely differ among local populations (Voje et al., 2013). It is noteworthy in this respect that our results point to evolutionary increases in allometry slope occurring in the deepest branches of our analysis, as the northern and southern lineages of *Trypoxylus* diverged from their shared common ancestor with *X. davidis* – a split estimated to have occurred almost 30 million years ago (Jin et al., 2016) – while all of the evolutionary changes to horn allometry that we observe among populations of *Trypoxylus* involve shifts in allometry intercept rather than slope (Figures 3, 4).

In the context of animal communication, increases in static allometry slope are predicted for strongly sexually selected structures (Alatalo et al., 1988; Fitzpatrick 1997; Kodric-Brown et al., 2006; Cuervo and Møller 2009; Knell et al., 2013; Hone et al., 2016), particularly those that function as conspicuous signals in agonistic assessment or mate choice (Fromhage and Kokko 2014; Voje 2016; Eberhard et al., 2018; O’Brien et al., 2018; Rodriguez and Eberhard 2019).

Specifically, a steeper static allometry slope increases the range of among-male variation in trait size (hypervariability), amplifying otherwise-subtle differences in underlying male condition so that they are easier for receivers to see (Hasson 1991; Wallace 1987; Cotton et al., 2004b; Bonduriansky 2007; Bradbury and Vehrencamp 2011; Tazzyman et al., 2014).

The evolution of steep static allometry often results from increases in the developmental plasticity or nutrition-sensitivity of trait growth (Shingleton et al., 2008; Tang et al., 2011; Emlen et al., 2012; Warren et al., 2013; Okada et al., 2019), and this ‘heightened’ condition-sensitive expression is yet another characteristic common to conspicuous structures that function as reliable signals of male body size or condition (Knell et al., 1999; David et al. 2000; Cotton et al. 2004a; Bonduriansky and Rowe 2005; Emlen et al. 2012; Warren et al. 2013). Indeed, such heightened conditional expression has already been demonstrated for the horns of this species (Emlen et al., 2012; Johns et al., 2014). Consequently, the dramatic increases in static allometry slope observed in the northern and southern lineages of *T. dichotomus* are consistent with animals beginning to use this structure as a signal, either by males to assess the resource holding potential of rival males, or by females.

Interestingly, there is no evidence that horns in this species actually function as a signal. Interactions among beetles take place at night, and although beetles do appear to be capable of discerning a structure like a horn (Obata and Hidaka 1983), all behavioral observations to date suggest that females ignore the horns entirely (Siva Jothy 1987; Hongo 2003, 2007, 2012; del Sol et al., 2020). Male fights unfold in a fashion consistent with some form of assessment during battles (e.g., fights are most likely to escalate when males are evenly matched [Hongo 2003, 2007; Karino et al., 2005; del Sol et al., 2020]). However, they do not posture or display the horn in any way conducive to visual assessment. It is possible that males use their horns as a tactile (rather than visual) signal during contests (Hongo 2003), as with the elongated forelegs of the flower beetle *Dicronocephalus wallichii* (Kojima and Lin 2019) and the elongated mandibles of *Cyclommatus mniszechi* stag beetles (Chen et al., 2022). Simply being able to touch an opponent first may suffice as a signaling function for structures like beetle horns, but this will need to be investigated in future studies.

Regardless of whether rhinoceros beetle horns function as a signal, they definitely are used as tools of battle. Males in all populations studied to date fight with rival males for feeding territories on the sides of host trees (Siva Jothy 1987; Hongo 2003, 2007, 2012; del Sol et al., 2020). Males attempt to position their head horn underneath the body of an opponent, lifting and flinging them away from the tree (McCullough et al., 2014; Buchalski et al., 2019), and males with the largest body sizes and longest horns consistently win these contests (Siva Jothy 1987; Hongo 2003, 2007; Karino et al., 2005; del Sol et al., 2020).

The two largest increases in male horn length were each accompanied by significant reductions to horn lifting strength (transitions A, E in Figures 3, 4 respectively), consistent with the ‘paradox of the weakening combatant’ (Levinton and Allen 2005). Then, as beetles colonized the northern Ryukyu islands of Kuchinoerabu-Yakushima-Tanegashima and Goto, male horns subsequently evolved to be relatively shorter, increasing their lifting strength in each instance. This may be the result of selection for horn strength offsetting the benefits of long horns, shifting the balance of selection on this structure. Alternatively, local changes to the breeding ecology of the beetles may have relaxed the strength of selection on horn length. Field observations of contemporary populations on Yakushima island suggest that beetle densities are much lower than on the mainland and feeding territories are more numerous, both of which could detract from the fitness benefits of long horns (Hongo 2006; del Sol et al., 2020; Kojima personal communication). On Goto island beetle densities are sometimes high and male fights and fight-related injuries are prevalent (Kojima personal observation), so the recent reduction in horn length could be interpreted as compensatory evolution to restore horn lifting strength.

In the southern lineage of *Trypoxylus* we see more compelling evidence for compensatory evolution. When beetles colonized Taiwan they retained the long horn lengths of their mainland Chinese neighbors, but these animals have at least partially ameliorated the associated reduction in horn lifting strength through an evolutionary increase in the height of their heads. Taller heads increase the length of the input lever (Figure 1), compensating for the long head horn and restoring strength to this exaggerated male weapon (Figure 4). Evolutionary elongation of stag beetle (Lucanidae) mandibles also appears to have been accompanied by compensatory increases in the length of the input lever (Goyens et al., 2016). The lever system of the stag beetle mandible rotates side-to-side, in contrast with the up-down lifting rotation of *T. dichotomus* horns, so increases in stag beetle input lever length resulted in wider, rather than taller, male heads, as well as substantially stronger mandible-squeezing forces (Goyens et al., 2016).

Here we assume that horn force production is important for winning fights and observations of both fighting behavior (Hongo 2006; del Sol et al., 2020) and beetle grip strength (McCullough et al., 2014) suggest that this is likely true. However, there are alternative biomechanical interpretations of the variation in morphology that we see across populations. The function of the horn lever system in combat is complicated. Horns are used to prod, slowly pry, and rapidly flick opponents from several different orientations. It is possible that different biomechanical solutions are used with different details of the behavioral ecology of different populations. There should be a force versus speed tradeoff associated with horn length and a longer horn relative to body size moves faster (Levinton and Allen 2005), therefore if speed of movement is important for winning contests, a longer horn may be directly selected for. Additionally, an increase in the input lever length of the system should increase the amount of force produced for a given muscle cross section, as described above, or alternatively should decrease the amount of muscle needed to produce a given force. Therefore, the increased mechanical advantage seen in Taiwanese beetles could be an example of selection for increased force, or for decreased energetic cost for producing a given force.

Field studies of contemporary populations on Taiwan consistently observe high densities of beetles – the highest yet recorded for this species – and male battles are both intense and frequent (McCullough 2012; 2014; del Sol et al., 2020), as would be expected if selection for male fight performance were driving the evolution of mechanical compensation at this location. Because the head height (input lever) of beetles from Okinawa/Kumejima and mainland China have similar allometries with the outgroup species (Figure S7), we are able to deduce the sequence of trait evolution in this clade: an elongated output lever (increase in horn length; Transition E) evolved first, followed by a subsequent increase in the input lever (head height; Transition F).

This provides one of the clearest illustrations to date of the evolution of mechanical compensation in a sexually selected male weapon system.

Recent research on four bar linkage systems suggests that components of mechanical systems with high mechanical sensitivity should evolve faster than the other parts of the system (Muñoz et al., 2018). If this prediction holds for the rhinoceros beetle horn lifting system, then the output lever (horn length) should be under selection for performance during fights (either lifting speed or force), resulting in evolutionary changes in system components with especially high mechanical sensitivity. However, the horn may also function as an agonistic deterrent signal, in which case the size of the horn *per se* may be under selection independent of mechanical performance. It will be interesting to compare rates of evolution in horn length, muscle area, and input lever length, to understand how mechanical sensitivity and direct selection on weapon size might interact to shape patterns of evolution of extreme, multi-function structures like rhinoceros beetle horns.

## Supporting information

R stats script

## Acknowledgments

This project was funded by NSF IOS–1456133 (DJE), JSPS 17H06901 (WK), The Gonzaga Science Research Program (BOS), NSF IOS-1456731 (LCL), MOST 103-2311-B-029-001-MY3, 104-2621-B-003-002-MY3 (C-PL). Benjamin Buchalski and Dylan Scanes helped collect biomechanical measurements.

